# Cystoliths in *Ficus* leaves: increasing carbon fixation in saturating light by light scattering off a mineral substrate

**DOI:** 10.1101/2020.04.08.030999

**Authors:** Maria Pierantoni, Indira Paudel, Batel Rephael, Ron Tenne, Vlad Brumfeld, Shai Slomka, Dan Oron, Lia Addadi, Steve Weiner, Tamir Klein

## Abstract

The manner in which leaves adapt to different light intensities is key for enabling plants to survive in diverse environments and in constantly changing conditions. Many studies have addressed this subject, but little attention has been given to the effect that mineral deposits in leaves can have on photosynthesis.

Here we study 6 species of *Ficus* and investigate how different cystolith configurations affect photosynthesis in both non-saturating and saturating light. We quantified the effect of light scattering by cystoliths on light absorption by measuring chlorophyll fluorescence intensity using microfluorimetry. We complement this by carbon assimilation measurements to directly estimate how light scattering by cystoliths affects the overall photosynthetic process.

We show that light waste is reduced when irradiance is on a cystolith compared to cystolith free tissue. Moreover, light is channeled into the center of the leaf where photosynthesis occurs more efficiently than in the outer layers. This, in turn, leads to more efficient CO_2_ assimilation.

We conclude that cystoliths contribute to photosynthesis optimization under saturating light. Cystoliths reduce the wasted portion of absorbed light under saturating irradiance by scattering light into the light-deprived leaf center. The increased efficiency may well provide important benefits to plants that form mineral scatterers.

## INTRODUCTION

Leaves are adapted to collect photons and use them for photosynthesis. When light is dim, the low amount of photons reaching the leaf limits photosynthesis. However, under most irradiance regimes photons are not the limiting factor for photosynthesis, as only a small portion of the light absorbed by the leaf is used for photosynthesis. The remaining light is dissipated either in the form of chlorophyll fluorescence or as heat (i.e., photoprotection) (Adams III & Demmig-Adams 1993, Niyogi 2000, Powles 1984). Little is known about the strategies evolved by leaves to reduce the proportion of wasted photons and, subsequently, to enhance photosynthesis (Smith et al. 1997, Terashima & Hikosaka 1995, Wittenberg et al. 2014).

The palisade cells below the adaxial (upper) surface of the leaf are specialized for photon collection, and in many leaves these cells are responsible for most of the light harvesting. Leaves can also absorb scattered light and collect photons in the lower mesophyll below the abaxial surface (Vogelmann 1989). In fact, although the upper, palisade tissue, is more efficient in light absorption than the lower, spongy tissue, the latter has higher light absorption per unit photosynthetic pigments (Xiao et al. 2016). However, a major difficulty for the leaf to efficiently use the incident photons on both surfaces is that the uppermost layer of cells absorbs almost all the light, creating a steep light gradient from the leaf surface to the leaf interior (Vogelmann 1993, Vogelman et al. 1996, Zhu et al. 2010, Pierantoni et al. 2018). Thus, under most irradiance regimes the impinging light exceeds the saturation threshold (Vogelmann et al. 1989, Ort 2001, Sušila et al. 2004). Recently, a leaf ray tracing model showed how chloroplast positioning within leaf cells and the leaf internal light environment affect the leaf light-use efficiency (Xiao et al. 2016). Light sheet microscopy showed that light attenuation was more gradual in a low-chlorophyll mutant soybean (Slattery et al. 2016). Photoprotection dissipates the unused photons to prevent light-induced damage (Powles 1984, Adams III & Demmig-Adams 1993, Havaux & Niyogi 1999, Niyogi 2000, Niyogi 2017). When photoprotection occurs, CO_2_ assimilation becomes the rate-limiting factor for the photosynthetic process (Von Caemmerer & Farquhar 1981, Niyogi et al. 1997). The maximum rates of CO_2_ fixation occur in the middle of the leaf (Nishio et al. 1993). Consequently, increasing the irradiance in the center of the leaf and reducing the excess light on the uppermost leaf layers can be a major advantage in terms of carbon fixation efficiency.

Anatomical strategies to increase light penetration into the lower mesophyll include, e.g., bundle sheath extension in heterobaric leaves (Nikolopoulos et al. 2002). Other leaves have evolved a strategy that uses mineral scatterers to introduce light deep in the leaf tissue and by so doing enhance photosynthesis. This possibility was first proposed by Kuo-Huang et al. (2007), and documentation showing that mineral scatterers channel light deep into the tissue and use it for photosynthesis, was provided for *Carya* and *Ficus* trees by Gal et al (2012). The effect of mineral scatterers on photosynthesis was measured by microfluorimetry, using both the transient chlorophyll fluorescence response upon exposure to saturating light and the overall chlorophyll fluorescence intensity immediately following exposure (Gal et al. 2012). Notably, both of these properties were found to be highly correlated with one another, reporting on the wasted portion of absorbed light and therefore serving as an inverse proxy for the efficiency of photosynthesis (Govindjee 1986, Papageorgiou 2007). Measurement of chlorophyll fluorescence has the advantage of accessing local differences in fluorescence with micrometer resolution, and was used in two follow-up studies on various *Ficus* species (Pierantoni et al 2017, 2018). Microfluorimetry can thus be used to differentiate between the effects of irradiance on minerals and on the surrounding tissue. Microfluorimetry cannot, however, directly measure the yield of the photosynthetic process.

Direct measurements of changes in carbon fixation deriving from photosynthesis can be obtained by monitoring CO_2_ or O_2_ evolution (Bjorkman 1971, Von Caemmerer & Farquhar 1981). To date, a direct measurement of the effect of light scattering by minerals on photosynthesis was never performed. As carbon assimilation measurements are not performed at micrometer resolution, distinguishing the effect of minerals from the rest of the tissue is challenging. Ideally, the effect of mineral bodies on photosynthesis should be studied on the same plant species with and without minerals. Unfortunately, to our knowledge, such plants are not available. Furthermore, mineralization cannot be inhibited without affecting the health of the whole plant.

In this study we use a different approach to assess the direct effect of *Ficus* leaf minerals on photosynthesis. *Ficus* species can inhabit very different environments, some of which are characterized by very strong irradiance (Janzen 1979, Harrison 2005, Zhang et al. 2016), and some *Ficus* species are known to have among the highest rates of photosynthesis in plants (Zotz et al. 1995, Hao et al. 2011). Leaves from many *Ficus* trees deposit hydrated amorphous calcium carbonate bodies, called cystoliths, just below the leaf surfaces (Meyen 1839, Ajello 1941, Omori & Watabe 1980, Setoguchi et al. 1989). Here we study leaves from six different *Ficus* species *(F. religiosa, F. bengalensis, F. microcarpa, F. benjamina, F. lyrata, F. carica)*. In these species cystoliths are deposited adaxially (below the upper surface of the leaf), abaxially (below the lower surface of the leaf), on either sides, or are not deposited at all. We hypothesize that the presence and distribution of cystoliths within the leaf tissue is correlated with variations in photosynthesis (higher CO_2_ assimilation) and photosynthetic efficiency (lower chlorophyll fluorescence). By comparing the different species under both non-saturating and saturating illumination conditions, we can deduce the direct effect of adaxial and abaxial cystoliths on carbon fixation and study their dependence on the light regime.

## MATERIALS AND METHODS

### Plant Material

Our study was entirely conducted on the Weizmann Institute of Science campus (Rehovot, Israel). The campus gardens have a large collection of tree species of various biomes (tropical, subtropical, Mediterranean, temperate, etc.), including six *Ficus* species (*F. religiosa, F. bengalensis, F. microcarpa, F. benjamina, F. lyrata, F. carica*). Single mature trees growing outdoors were used. All the tree species were within 1 km. Mature leaves from *Ficus* trees (*n* = 6 per species) were freshly collected and immediately scanned. For gas exchange measurements, the same mature trees were measured, along with saplings of the same *Ficus* species, which were purchased from a local nursery and grown at the Weizmann greenhouse facility during November 2017. Three saplings of each species were grown under controlled conditions for at least 2 weeks (22-25° C daytime temperature and ~50% relative humidity). Saplings were at 1.0 ± 0.3 m height, except *F. carica and F. benjamina*, which were 1.5 ± 0.4 m. Stem base diameters were 3.5 ± 0.5 cm across all saplings. Among the six species, five were evergreen, with *F. carica* being the only winter deciduous. Nevertheless, under the warm greenhouse conditions and the *in situ* thermo-Mediterranean climate, both *F. carica* saplings and mature trees maintained leaves during measurement days, which were mostly between June 2017 and December 2018.

### Micro-Computed Tomography (microCT)

We applied X-ray micro-computed tomography, an increasingly useful tool in tree anatomy and physiology (e.g., Choat et al. 2016), to detect the presence and localization of cystoliths inside leaves. We preferred to use MicroCT over other anatomical methods, because the cystoliths, being the object of investigation, are denser than the surrounding leaf environment, and are hence easily detectable with the X-ray source. MicroCT scans were acquired using a Micro XCT-400 (Zeiss X-ray Microscopy, California, USA). Triangular sections of leaves (base 1 cm, length 2 cm) were cut and placed in a sealed plastic pipette tips. To prevent dehydration, the tips were partially filled with water, leaving part of the leaf in air. Only the part of the leaf not immersed in water was imaged. The tomographic image was obtained by taking 1300 projections (180 degrees) at 40 kV and 800 μA. The final voxel size was 1.5 μm. 3D volumes were produced using the Avizo 3D analysis software (FEI Visualization Sciences Group, Berlin, Germany). Leaf thickness, cystolith volumes, percent of leaf surface covered by cystoliths, cystolith dimensions were obtained using the same software. Minerals that are the most absorbing bodies in the leaf were selected by contrast thresholding. The leaf total volume was calculated by manual segmentation and interpolation of the selected volume segments. The reproducibility of the values was estimated by repeating the measurements three times. A variance of 0.25% was obtained.

### Micro-modulated Fluorimetry

Measurements were performed on freshly sampled leaves, immediately after detachment. To perform leaf auto-fluorescence measurements a micro-fluorimeter setup was custom built around a commercial microscope (Eclipse Ti-U, Nikon) as in Pierantoni *et al.* (2018). A 635 nm pulsed diode laser (EPL635, Edinburgh Instruments) emitting pulses with a 100ps duration at a 20 MHz repetition rate excited the chlorophyll in the leaf. The beam was focused through a 250mm focal length lens (LA1461, Thorlabs) onto the back aperture of an objective lens (20X, 0.4 N.A., Nikon). As a result, a ≈12 μm diameter spot was formed in the sample plane. Control over the laser excitation power was achieved by using a variable retarder between two cross-polarized linear polarizers and a mechanical shutter was used to start and stop irradiance periods. Fluorescence light was collected through the same objective, filtered from the direct irradiance of the laser by a dichroic mirror (650LP Semrock) and a dielectric filter (635LP Semrock), and imaged onto an electron-multiplying charge-coupled device (EMCCD) camera (iXon Ultra 897, Andor). The camera captured a 50 s long frame series with a ≈20 ms exposure time to accurately capture the fluorescence dynamics. Dynamic traces started with 5s without any excitation followed by a dim excitation period (typically 1 nW, matching a flux of 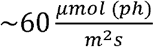) with a 5s duration in order to obtain the photoluminescence quantum yield for unsaturated reaction centers, followed by a 10 s dark period, followed by another period (30 s) of saturating excitation power (Fig. S1; typically 15 nW, matching a flux of 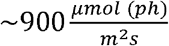). Integrated fluorescence intensities were calculated by summing the EMCCD signal over the entire saturating period (30 s) on a ~80 *μm*^2^ region of interest (Fig. S1). Data for *F. microcarpa* and *F. carica* were taken from Pierantoni *et al.* (2018). The data were recorded and analyzed with a custom written MATLAB script. Comparisons between fluorescence intensities in C locations and T locations were performed using a Student *t*-*test* (two tailed, homoscedastic) with significance level of 0.05 both for adaxial and abaxial measurements.

### CO_2_ Assimilation and Stomatal Conductance

Leaf CO_2_ and H_2_O gas exchange measurements were carried out in young, mature, leaves using the infra-red gas analyzer Walz GFS-3000 photosynthesis system (Walz, Effeltrich, Germany). The ambient CO_2_ concentration was 400 ppm (setup reference CO_2_ levels using external sources), leaf temperature was fixed to 25 °C and flow rate was 750 μmol s^−1^. All measurements were performed between 9:00 h and 12:00 h during June 2017-February 2018. In the greenhouse all the measurements were conducted on intact, attached leaves. Due to tree height, measurements on mature trees were performed after detaching small twigs with about 5-10 leaves which were exposed to full or partial sunlight for at least 30 minutes. Leaf gas exchange remained unchanged ~15 minutes following detachment, and hence our measurements, which were always performed within 3-8 minutes, captured the native state of the leaf. Thanks to the relatively large leaf size across *Ficus* species, measurements were performed on 8 cm^2^ sections in the middle of each leaf. Sensors of the infra-red gas analyzer were calibrated according to manufacturer’s guidelines and zeroed prior to each measurement. Each leaf section was illuminated firstly adaxially keeping the abaxial side covered, and then abaxially keeping the adaxial side covered, ~5 min each. Experiments were conducted using both saturating irradiance (1000 μmol (photon) m^−2^s^−1^) on sunlit leaves, and dim, non-saturating irradiance (500 μmol (photon) m^−2^s^−1^) on partly shaded leaves. Leaf selection was intended to make sure that leaves were acclimated to the light intensity at the time of measurement. Stomatal aperture can be directly measured only by observation of each leaf species under the microscope. For leaves belonging to the same species, where stomatal density and stomatal size are homogeneous, stomatal aperture is directly proportional to stomatal conductance (Klein 2014, Bartlett et al. 2016). Stomatal conductance is the ratio of leaf transpiration (the process by which gas molecules move through the stomata) and vapor pressure deficit (the difference in vapor pressure between the air and the leaf). Both parameters were measured directly by the Walz photosynthesis system, and stomatal conductance was readily calculated. Pairwise comparisons between net photosynthetic rate in adaxial side and abaxial side (Pn_adaxial_ and Pn_abaxial_) acquired in saturating light and dim light for outside trees and greenhouse sampling were performed using a Student *t*-*test* (two tailed, paired) with significance level of 0.05.

## RESULTS

### Cystoliths and Cystolith Organization in the Leaf Tissue

We studied leaves from six species of *Ficus* that have diverse patterns of cystolith types and distributions (Fig. 1). MicroCT was applied, exposing the cystoliths embedded within the leaves, thanks to their higher density relative to the leaf tissue. *F. religiosa* leaves do not have cystoliths (Fig. 1a). *F. bengalensis* leaves (Fig. 1b) deposit adaxial cystoliths whose morphologies and sizes vary (Table 1). Some of the cystoliths do not extend into the palisade, but rather grow in the epidermis, i.e. they are oriented perpendicular to the palisade layer (Fig. S2 a). These *F. bengalensis* cystoliths are hollow in the center, and their distribution is less organized than in the other species. The leaves of *F. bengalensis* represent the only studied case where cystoliths are deposited also above the veins (Fig S2). *F. microcarpa* and *F. benjamina* are similar in that both species deposit cystoliths both adaxially and abaxially at regular distances (Fig. 1c, d). The adaxial cystoliths are all elongated and extend into the palisade, whereas the abaxial cystoliths are smaller and more rounded. In *F. microcarpa* the adaxial cystoliths are larger than in *F. benjamina* (Table 1), whereas the abaxial minerals are comparable in size in the two species (Table 1). *Ficus lyrata* deposits elongated cystoliths on the adaxial side, forming a regular pattern (Fig. 1e). Some of the mature leaves contain very few abaxial cystoliths. In *F. carica*, spherical cystoliths are deposited on the abaxial side, while adaxial cystoliths are absent (Fig. 1f). *F. carica* is the only species among the studied species, to deposit abundant silica in the epidermis and in the hairs (silicified hairs are visible in Fig. 1f). All six *Ficus* species deposit calcium oxalate minerals along their veins.

**Fig. 1.**
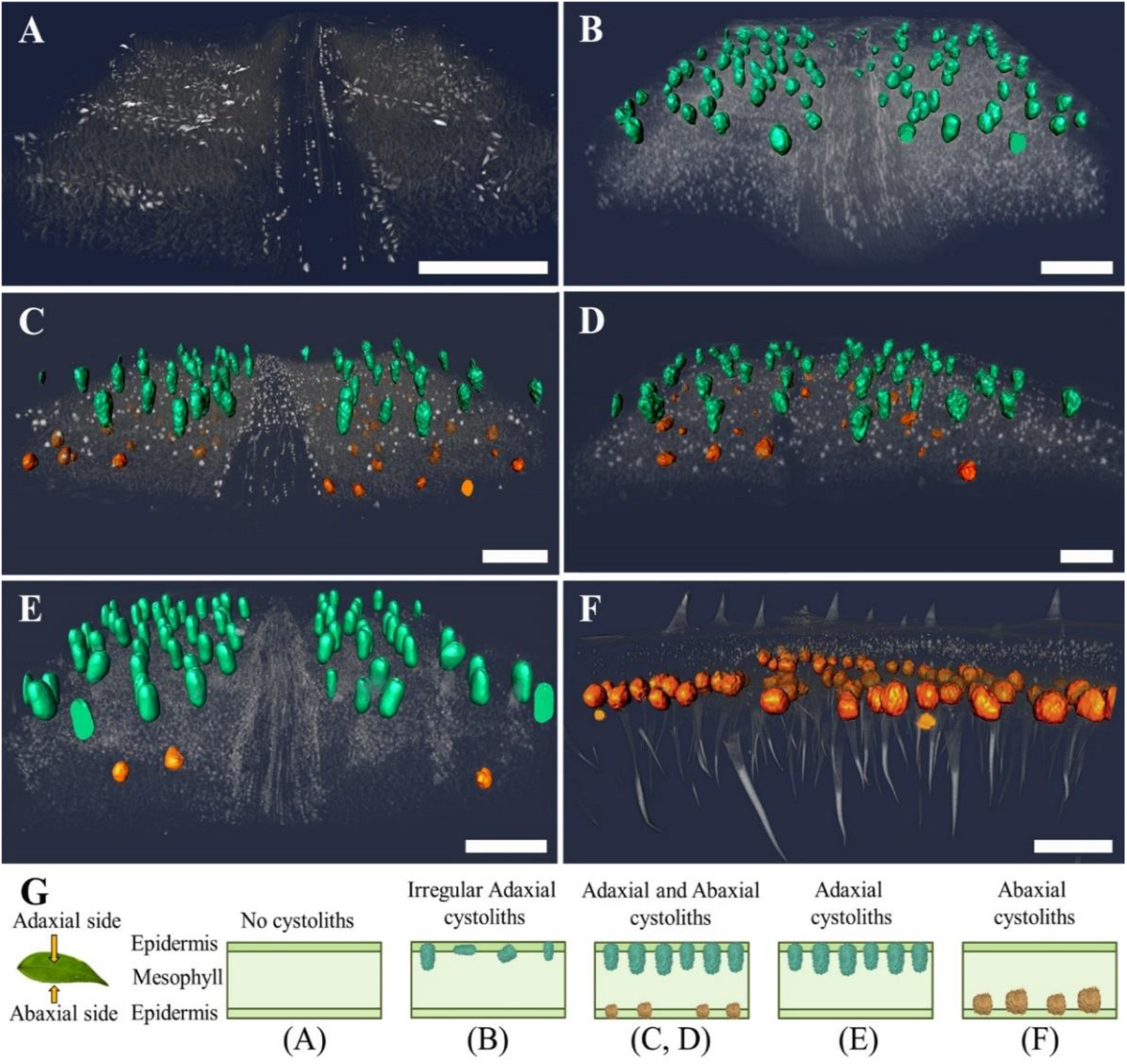
MicroCT 3D cross-sectional perspective views of minerals and soft tissues in *Ficus* leaves. The cystoliths are artificially colored. Adaxial cystoliths are colored in cyan and abaxial cystoliths in orange. The dark grey colored textures are the soft tissues and the lighter grey are the relatively small calcium oxalate crystals. In *F. carica* silicified hairs are also visible. All leaves are shown with their upper adaxial surfaces on the top. (A) *F. religiosa*, (B) *F. bengalensis*, (C) *F. microcarpa*, (D) *F. benjamina*, (E) *F. lyrata*, (F) *F. carica*. Scale bars: 300 μm. (G) Schematic representations of the different *Ficus* leaf cross sections showing cystolith locations and morphologies in the soft tissue.

**Table 1.** Leaf thickness, cystolith volumes, percent of leaf surface covered by cystoliths, cystolith dimensions in the *Ficus* leaves investigated and corresponding standard errors. Leaf parameters were calculated from 6 different leaves for each species, cystolith sizes and stomatal parameters were all obtained for n=50.

| Leaf | Leaf thickness (µm) | Cystolith volume content | ADAXIAL SIDE (µm) |  |  |  |  | ABAXIAL SIDE (µm) |  |  |  |  |
| --- | --- | --- | --- | --- | --- | --- | --- | --- | --- | --- | --- | --- |
|  |  |  | Leaf area covered by adaxial cystoliths | epidermis | stomata | palisade | cystoliths | Leaf area covered by abaxial cystoliths | lower mesophyll | cystoliths | epidermis | Stomata (inter-stomatal distance) |
| *F. religiosa* | 210±40 | / | / | 15±2 | / | 70±10 | / | / | 110±20 | / | 15±4 | 15±5<br>(45±30) |
| *F. microcarpa* | 320±20 | 1.3±0.05% | 4.5±0.5% | 45±10 | / | 85±20 | 100±20 | 2.4±0.2% | 155±5 | 45±10 | 35±10 | 20±4<br>(50±20) |
| *F. benjamina* | 220±10 | 1.4±0.08% | 4.4±1% | 10±2 | / | 75±10 | 90±15 | 1.2±0.6% | 110±20 | 45±10 | 15±5 | 18±6<br>(30±5) |
| *F. bengalensis* | 340±20 | 1.6±0.3% | 8±1.2% | 45±20 | / | 115±20 | 60±30 | / | 160±20 | / | 20±10 | 16±3<br>(18±7) |
| *F. lyrata* | 440±20 | 2.3±0.1% | 7±0.5% | 40±15 | / | 120±5 | 100±15 | / | 250±10 | 40±20 | 30±15 | 10±2<br>(20±4) |
| *F. carica* | 240 | 1.6±0.2% | / | 30±5 | / | 100±10 | / | 8.9±1.6% | 100±10 | 80±20 | 10±5 | 15±3<br>(10±5) |

The volume of the leaf occupied by cystoliths varied from 1.3% in *F. microcarpa* to 2.3% in *F. lyrata*, while cystoliths cover from 1.2% of the leaf surface for *F. benjamina* abaxial cystoliths, to 8.9% of the leaf surface in *F. carica abaxial cystoliths* (Table 1). A schematic representation of all the cystolith deposition patterns is presented in Fig. 1g. In general, across all the *Ficus* species examined here, the palisade cells were compact and elongated, while the lower mesophyll is formed by round and separated cells (Fig. S3). Nevertheless, each species has its own soft tissue organization. In *F. religiosa*, palisade cells are less compact than in other species, with air spaces between the lower mesophyll cells (Fig. S3 a). In *F. lyrata*, the upper palisade is formed by very compact cells (Fig. S3 b). The lower mesophyll layer is twice as thick as the palisade and is formed by small spherical cells (Fig. S3 b). In *F. carica* the palisade and lower mesophyll are of similar thickness (Fig. 2c). The *F. carica* palisade cells are elongated and close to each other, whereas the cells in the lower mesophyll are more rounded and more spaced (Fig. S3 c). *F. microcarpa*, *F. benjamina* and *F. bengalensis* have a very similar soft tissue structure to *F. lyrata* but in these three species, the lower mesophyll is just slightly thicker than the palisade.

**Fig. 2.**
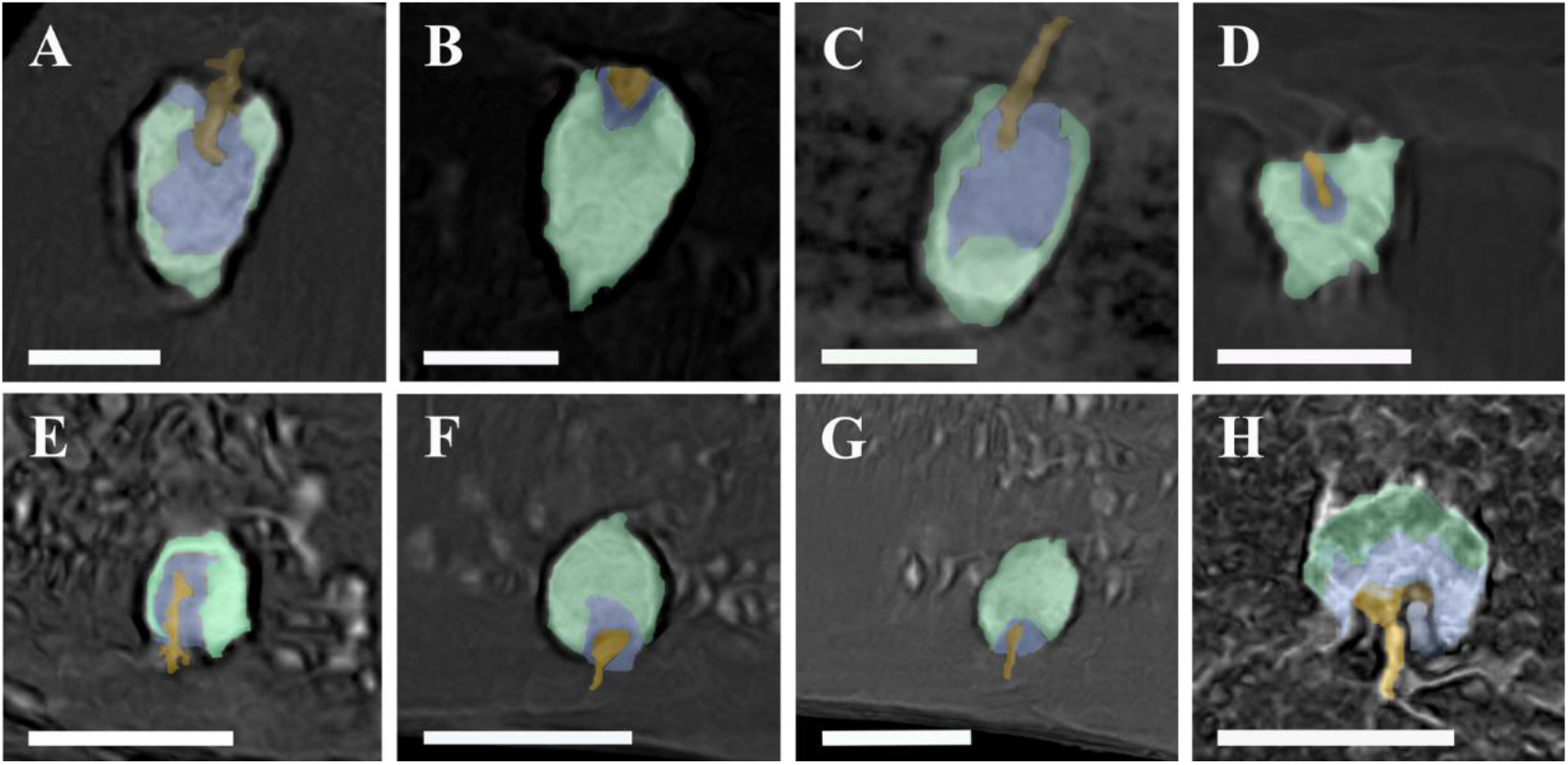
MicroCT sections showing that the cystoliths studied all have the same basic composite structure. (A-D) adaxial cystoliths; (E-H) abaxial cystoliths. (A, E) *F. microcarpa*; (B, F) *F. benjamina*; (C, G) *F lyrata*; (D) *F. benghalensis*; (H) *F. carica*. yellow = silica stalk; violet = internal amorphous calcium carbonate (ACC) phase; green = external ACC phase. Scale bars = 60 μm.

### Chlorophyll Fluorescence

Microfluorimetry measurements were performed on the six *Ficus* species. Cystoliths and control areas of tissue without cystoliths (C=cystolith and T=tissue locations respectively) were identified by transmitted light microscopy. We note that in a previous study (Gal et al. 2012), the local fluorescence intensity measurement was found to be highly correlated with changes in the time constant of the transition from lower to higher fluorescence levels, and were shown to be a good proxy of the amount of the degree of saturation and the amount of wasted light. Laser irradiance was performed in C and T locations both adaxially (Fig. 3 a) and abaxially (Fig. 3 b). Where minerals were absent on one of the leaf sides, only T locations were measured. The irradiance intensity was 900 μmol (photon) m^−2^s^−1^ within the absorption band of Chlorophyll b, an insolation regime characteristic of the tree growth conditions (Fig S1). Each bar of the histogram in Fig. 3 is the value obtained by averaging 12-26 measurements (number of measurements indicated under each bar) from C and T. For adaxial measurements when the irradiance was on cystoliths (C locations) the wasted light was reduced by (41±20)-(76±15)% compared to T locations (Fig. 3). For abaxial measurements, in the cases of *F. microcarpa* and *F. benjamina*, cystoliths reduced wasted light by (72±25)% and (55±30)%, respectively (Fig. 3). For *F. carica* abaxial surfaces, there was no difference in fluorescence between C and T locations. Overall, cystoliths always improved light harvesting for adaxial irradiance. *F. microcarpa* and *F. benjamina* improved light harvesting also when light reached the abaxial leaf side. A student t-test was conducted to determine the significance of the observed difference between fluorescence intensities in C locations and T locations, both for adaxial and abaxial measurements (Fig. 3). Fluorescence was significantly different for C and T locations across the species, apart for *F. carica.* This might be due to the fact that in *F. carica* the dense silica hairs prevented precise localization of the cystoliths, in turn scattering both incident and emitted light. For all the leaves, except those of *F. carica*, the average fluorescence for adaxial T locations (Fig. 3a, light gray bars) was lower than for abaxial T locations (Fig. 3b light gray bars). Therefore, light harvesting was higher for adaxial irradiance than for abaxial irradiance. In *F. carica*, adaxial and abaxial fluorescence intensities were comparable. The same trend is obtained if, instead of comparing the average fluorescence intensity for the 6 *Ficus* species, single maximum fluorescence intensity values are considered (Fig. S6).

**Fig. 3.**
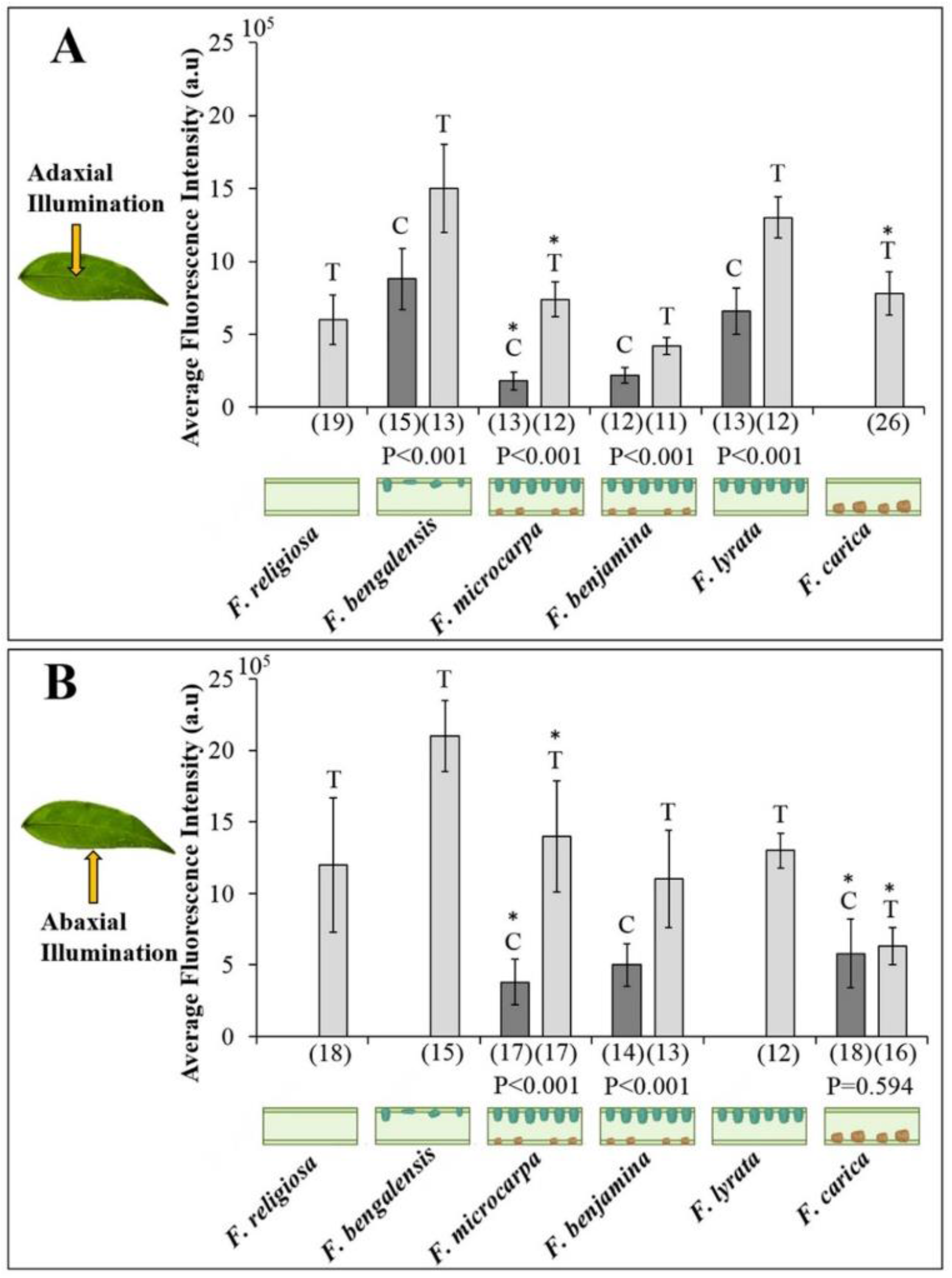
Bar plots of average fluorescence intensity from microfluorimetry curves for the 6 *Ficus* species. The data were obtained illuminating (A) adaxially and (B) abaxially. Dark gray bars are for measurements on cystoliths (C) and light gray bars are for measurements on tissue without cystoliths (T). The number of measurements from which the average was calculated is indicated under the bar. P-values from Student t-test comparing fluorescence intensities in C locations and T locations are also indicated under the bar. Values in bold indicate significant difference at α = 0.05. A clear significant difference between irradiance on C and T sites is demonstrated in all cases but one. The standard error for each average is indicated by the error bar. Leaf cross sections showing cystolith locations and morphologies are schematically represented on the x axes. The data marked by the asterisks are taken from Pierantoni et al. 2018.

### Leaf CO_2_ Assimilation

We measured CO_2_ assimilation in order to directly quantify the effect of cystoliths on carbon fixation in photosynthesis. Net CO_2_ assimilation (Pn) was measured using adaxial and abaxial irradiance separately, at both 1000 μmol (photon) m^−2^ s^−1^ (comparable to midday in the sun) and 500 μmol (photon) m^−2^ s^−1^ (comparable to light intensity in the shade). For each leaf, the percentage variation between Pn for adaxial irradiance (Pn _adaxial_) and Pn for abaxial irradiance (Pn _abaxial_) was calculated as:

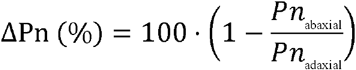

Discrete measurements are presented in Table S1. In general, palisade cells are compact and elongated light pipes that allow more efficient light harvesting than the round and spaced cells of the lower mesophyll (Fig. 2). Consequently, even when no cystoliths were formed, as is the case for *F. religiosa*, CO_2_ fixation was higher for adaxial than abaxial irradiance (positive ΔPn (%) values in Table S1). At 1000 μmol (photon) m^−2^ s^−1^ irradiance for *F. bengalensis, F. microcarpa, F. benjamina* and *F. lyrata* ΔPn (%) was higher than for *F. religiosa*. For *F. carica* ΔPn (%) was negative, meaning that the photosynthesis was higher for abaxial irradiance (Fig. 4 and Table S2).

**Fig. 4.**
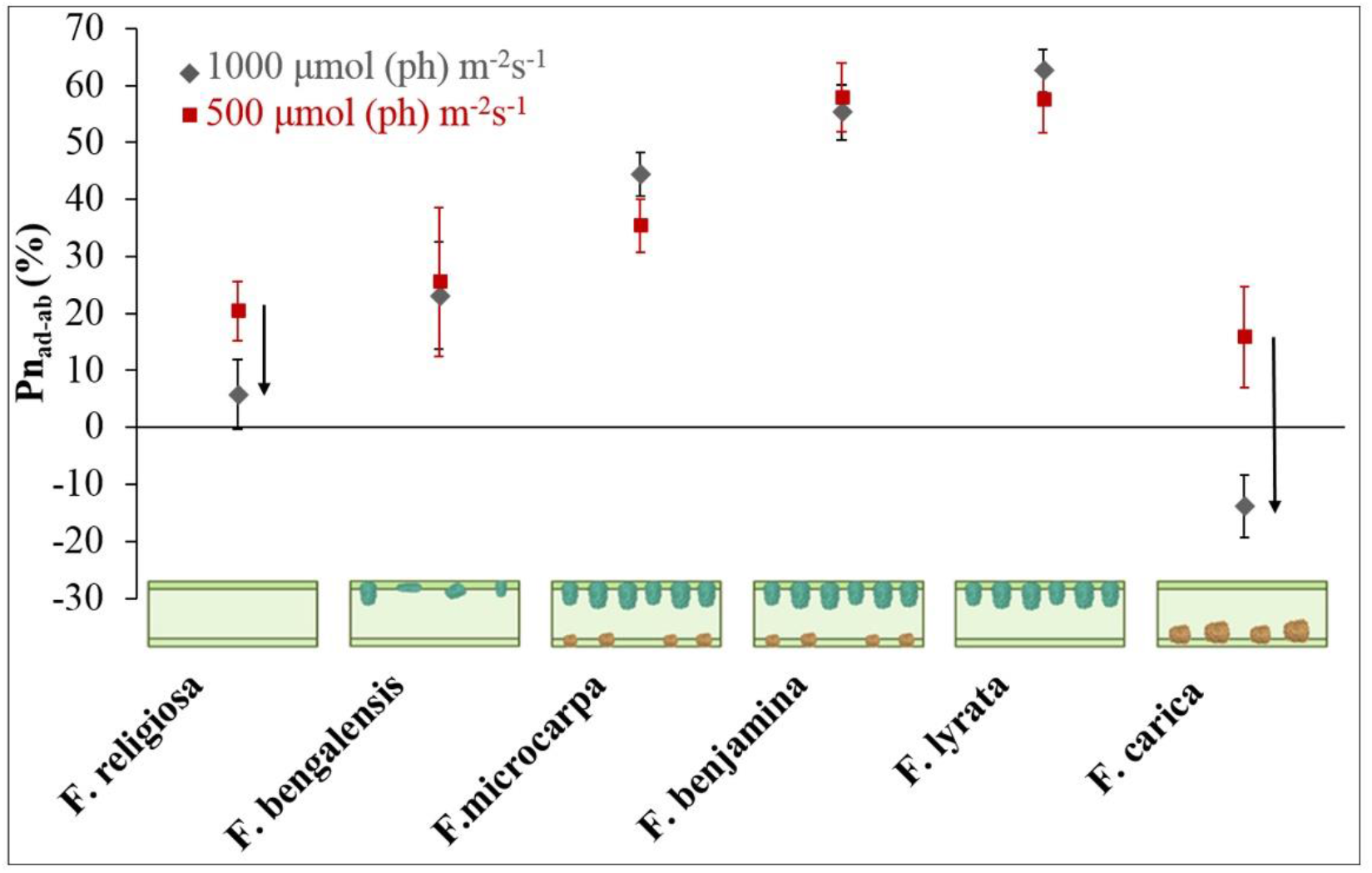
Average percentage variation between net CO_2_ assimilation for adaxial irradiance, Pn_ad_, and for adaxial irradiance, Pn_ab_, (ΔPn (%)). Dark gray diamonds: data obtained at 1000 μmol (ph) m^−2^s^−1^ irradiance. Red squares: data obtained at 500 μmol (ph) m^−2^s^−1^ irradiance. The bar indicates the standard error. The measurements were performed both on trees growing outdoor and in a greenhouse. In saturating or quasi-saturating light ΔPn (%) decreases only in the case of *F. religiosa* and *F. carica* (arrows).

Interspecific differences in the adaxial-abaxial percentage difference in the CO_2_ assimilation were smaller in high vs. low irradiance. Moving from a low light regime to high light, ΔPn (%) decreased in the case of *F. religiosa* and *F. carica* (Fig. 4, arrows), whereas for all other leaves ΔPn (%) did not change (*F. bengalensis, F. benjamina* and *F. lyrata* in Fig. 4) or even slightly increased (*F. microcarpa* in Fig. 4). Under low light, the *F. religiosa* palisade was not saturated but the lower mesophyll might have already been unable to use all the absorbed light. Thus, the intrinsic differences between the palisade and the lower mesophyll are important (ΔPn (%) = 20 ± 5). When the light was 1000 μmol (photon) m^−2^s^−1^ for both palisade and lower mesophyll, the difference in adaxial and abaxial net CO_2_ uptake decreased (ΔPn (%) = 6 ± 7). It is in this light regime that we expect mineral light scattering to be most important, by reducing the negative effect of saturation (or quasi-saturation) on photosynthesis.

The trend observed for ΔPn (%) was also obtained measuring quantum yield at the whole leaf scale. Quantum yield was higher for adaxial irradiance than for abaxial irradiance for the species depositing adaxial cystoliths (*F. microcarpa, F. benjamina*, and *F. lyrata*) compared to *F. religiosa*. In order to separate the contributions of the cystoliths from those of the soft tissue organization for each species, we subtracted from mean ΔPn (%) the value measured for *F. religiosa*: ΔPn_cyst_ (%) = ΔPn _*F.sp*_ (%) − ΔPn _*F.religiosa*_ (%). This operation was performed individually for the two irradiance regimes (Fig. S7). A positive ΔPn_cyst_ (%) value indicates that the ΔPn (%) was higher relative to that of a leaf without cystoliths. For values equal to or very close to 0, the difference in photosynthesis was probably solely due to the soft tissue. A negative ΔPn_cyst_ (%) value indicates that the photosynthesis during abaxial irradiance increased compared to the case in which no cystoliths are deposited. This only occurred for the *F. carica* under the high irradiance regime.

To test the cystolith effect on photosynthesis under a range of environmental conditions and tree ages, we compared leaves from mature, outdoor growing trees, with leaves from saplings growing in a greenhouse under controlled conditions (Fig. S8). ΔPn (%) was averaged for each species. Even if the specific recorded values vary, the averaged ΔPn (%) were not significantly different between trees growing outdoors and trees growing in the greenhouse (Fig. S7), indicating a similar conserved leaf physiology, independent of tree age and environment. Photosynthetic rate is directly affected by stomatal aperture, which is triggered by light stimuli (Humble et al. 1970, Sharkey and Raschke 1981). Consequently, we had to rule out the possibility that the differences in ΔPn (%) were solely due to differences in stomatal aperture response. We compared the adaxial-abaxial percent variation in stomatal conductance to ΔPn (%). The specific results obtained for three of the species are shown in Fig. S9. Apart from a few individual leaves, the variations in stomatal conductance were consistently smaller than variations in CO_2_ assimilation rates. These data show that the variations in stomatal aperture cannot be the main factor responsible for the recorded adaxial-abaxial percent variations in carbon assimilation.

## DISCUSSION

This study demonstrates a clear association between leaf cystolith deposits and increased photosynthesis in saturating light among *Ficus* species. This, therefore, confirms our hypothesis. Environmental factors and stomatal aperture variations are not the main factors responsible for determining interspecific differences in photosynthetic rates in saturating light. Instead, the redistribution of light deep into the leaf by cystoliths can significantly improve photosynthetic efficiency. This effect was observed using two different techniques: (1) Chlorophyll fluorescence measurements showed that less light was wasted when scattered deep into the tissue by cystoliths; and (2) CO_2_ assimilation measurements indicated that when the light is saturating, light scattering by cystoliths significantly increases photosynthesis. The results obtained with these two techniques cannot be directly correlated, but they independently show that light scattering by cystoliths can improve photosynthetic efficiency. In fact, the amount of absorbed light is increased and the light is channeled into the center of the leaf where the rates of CO_2_ fixation are the highest and where photoinhibition is minimal (Nishio *et al.* 1993). Consequently, in these conditions even a small increase in light absorbance efficiency can translate into a much greater increase in photosynthesis efficiency.

In *F. bengalensis*, the adaxial cystoliths are not well developed (sometimes hollow) and their distribution is uneven. Here we showed that these cystoliths only slightly improve photosynthesis by scattering some of the incident light into the center of the leaf (Fig. S3). In *F. microcarpa* and *F. benjamina*, where the adaxial minerals are well organized, the effect of the cystoliths is higher than in *F. bengalensis* (Fig. S3). The difference in photosynthesis between adaxial and abaxial irradiance in *F. microcarpa* and *F. benjamina* was presumably reduced by the presence of small abaxial cystoliths which increased light harvesting also from the abaxial side. In *F. lyrata* only large adaxial cystoliths are deposited and the Pn_cyst_ (%) was the highest among the studied species. In *F. carica*, where the cystoliths are deposited abaxially, the Pn_cys_ was negative. The results show that in saturating light for these leaves abaxial irradiance becomes more favorable. The microfluorimetry measurements did not however show the effect of the cystoliths. This could be due to the fact that the *F. carica* abaxial epidermis is covered by silicified hairs, which complicate the differentiation between cystoliths and the rest of the leaf surface. The presence of the silicified hairs could have resulted in an incorrect assignment of C and T locations. We also note that *F microcarpa* and *F. benjamina* have very similar mineral morphologies and organization. We cannot explain the differences in photosynthetic efficiency for adaxial versus abaxial irradiance only in terms of cystoliths, and therefore assume that differences in soft tissue structure, and possibly stomatal conductance, are also involved. Overall, the photosynthetic parameters we measured in the *Ficus* leaves were similar with earlier studies on *F. benjamina*, *F. religiosa*, and other congeneric species (Hao et al. 2011, Zhang et al. 2016), which also highlighted interspecific differences in light energy dissipation. Potentially, our results might explain why *F. benjamina* had 90% of maximum net assimilation rate at 633 μmol m^−2^ s^−1^ and *F. religiosa* at 956 μmol m^−2^ s^−1^, in spite of the higher rate in the latter (Hao et al. 2011).

Can reabsorption of fluorescence by chlorophyll account for the lower fluorescence emission related to cystoliths? Clearly, the optical properties of the palisade and spongy tissues differ (Xiao et al. 2016). Yet, we argue that the light reflectance and transmittance effects are mostly in the visible light spectra, and hence, any reabsorption should have little effect. Studying abaxially red/non-red variegated leaves of *Begonia*, the hypothesis that red pigments internally reflect/scatter red light transmitted by the upper leaf surface back into the mesophyll (thereby enhancing photon capture in light-limited environments) was negated (Hughes et al. 2008). In non-saturating light, the mineral scattering possibly plays a smaller part in photosynthetic enhancement. We infer that when saturation is not reached, and photoprotection is not occurring, a higher proportion of the incident radiation is used by the first tissue layers and thus channeling light deep inside the leaf is of less importance. Note however that in our study we only measured the effects of constant saturating light and constant dim light. Yet in natural conditions the light intensity reaching each leaf can change even within milliseconds (Rascher & Nedbal 2006, Kaiser et al. 2014). Plants are exposed to sun flecks or transient shading. Furthermore as the irradiance angle changes rapidly along the day, leaf cells may pass from full sunlight to shade within a second (Werner et al. 2001, Wang et al. 2006). Plants have evolved ways to adapt to such constantly changing conditions, and photoprotection defends the leaf from sudden excess of light (Adams III & Demmig-Adams 1993, Niyogi 2000). However, photoprotection has a significant negative impact on CO_2_ assimilation (Ögren & Sjöström 1990, Werner et al. 2001). In fact, carbon fixation is limited until the readjustment of the photo-apparatus state is complete (Zhu et al. 2004). Because of this delay, the decrease in total carbon uptake due to photoprotection was estimated to be between 12% and 32%, depending on light intensity, light incident angle, temperature, and chilling tolerance of leaves (Zhu et al. 2004). This study does not account for other factors, such as wind and leaf movements. Therefore, the effect due to the slow recovery after photoprotection is potentially higher.

Understanding how plants adapt to changes in light intensity is crucial for understanding how plants efficiently adapt to different environments and how they survive in constantly changing conditions (Ort 2001, Golan, et al. 2006, Murchie & Niyogi 2011, Ollinger 2011, Klein et al. 2013, Ort et al. 2015). For example, a climbing plant that has to adjust to both shade and high light environments in a tropical forest, acclimates by changing leaf orientation (Feild et al. 2001). Leaf size, specific leaf area, and chlorophyll concentration were higher in bamboo growing in shaded vs. open habitats (Yang et al. 2014). Leaf hydraulic efficiency was also found to control maximum photosynthetic rate, through vein positioning, across multiple land plants (Brodribb et al. 2007). Yet, little attention was given to the important effect that mineral deposits in leaves can have on photosynthesis. For plants competing to survive in a light saturated environment, even a small increase in photosynthetic efficiency can provide a great advantage over plants that do not have this capability. We note that many different plants have mineral bodies close to their leaf surfaces (Kuo-Huang et al. 2007, Gal et al. 2012, Horner 2012, Horner et al. 2012, Pierantoni et al. 2017a, Pierantoni et al., 2017b) and thus the effect may be widespread.

## CONCLUSIONS

Measurements of chlorophyll fluorescence and CO_2_ assimilation of *Ficus* leaves show that cystoliths contribute significantly to photosynthesis optimization under saturating light. This can occur also when the cystoliths are deposited on the abaxial leaf side. By redistributing light, cystoliths increase light absorption and, above all, allow photosynthesis to occur deep in the leaf where CO_2_ assimilation is more efficient. By so doing cystoliths, reduce the negative effect of excess light and significantly improve CO_2_ assimilation

## Acknowledgements

We thank Prof. Assaf Gal for helpful discussions and Netta Varsano for help with image preparation. L.A. is the recipient of the Dorothy and Patrick Gorman Professorial Chair of Biological Ultrastructure and S.W. of the Dr. Trude Burchardt Professorial Chair of Structural Biology.

## Author contributions

Maria Pierantoni performed most of the experiments and analyzed the data. Indira Paudel, Batel Rephael, Ron Tenne and Shai Slomka helped perform the experiments and analyze the data. Vlad Brumfeld provided expertise in microCT. Dan Oron provided expertise in the optics, and together with Lia Addadi, Stephen Weiner and Tamir Klein conceived the original project and supervised the experiments. Maria Pierantoni and Tamir Klein wrote the original drafts and all authors participated in the writing.

## Data availability

All data used in this paper are included in the text, figures, and supplementary data.

## Legends for Supporting Tables and Figures

**Table S1.** CO_2_ assimilation measurements were performed on the six Ficus species. For each leaf net CO_2_ assimilation (Pn) was measured for adaxial irradiance (Pn adaxial) and for abaxial irradiance (Pn abaxial). The perceptual variation between Pn adaxial and Pn abaxial was calculated (Pn ab-ad% = 100-100*Pn_abaxial_/Pn_adaxial_). P-values from a paired t-test was obtained by testing the difference between Pn_adaxial_ and Pn _abaxial_ The values are indicated in the first column in between brackets. Significant difference at α = 0.05.

**Table S2.** Average percentage variation between Pn for adaxial irradiance and Pn for abaxial irradiance (Pn_ab/ad_ (%)) obtained using 1000 μmol m^−2^ s^−1^ and 500 μmol m^−2^ s^−1^ irradiance values.

**Fig. S1**. Fluorescence intensity evolution over time. The graph shows the dynamics of the measurement for the 6 *Ficus* species on the adaxial side, off crystals. During a 30 s measurement the fluorescence intensity does not varies significantly. The decrease in fluorescence signal is about 10% for all leaves accept *F.bengelansis* with a 40% decrease. The same trend is shown considering adaxial measurements on crystals. From the curves it is also possible to appreciate that the 900 μmol (photon) m^−2^s^−1^ irradiance is suturing all leaves.

**Fig. S2.** MicroCT volumes of *Ficus bengalensis* leaves. A) top view showing that some of the cystoliths located above the main vein (delimited by yellow lines) are hollow in the center and they are elongated perpendicular to direction of the palisade layer. B) cross-sectional perspective view showing that some of the cystoliths (red arrows) are deposited also above the veins (the main vein diameters is shown by a yellow circle).

**Fig. S3.** MicroCT cross section of *Ficus* leaves showing tissue anatomy. In the palisade and the lower mesophyll single cells are highlighted respectively in green and blue. The air spaces between lower mesophyll cells are purple. (A) *F. religiosa*, (B) *F. lyrata*, (C) *F. carica* . Scale bar 100 μm.

**Fig. S4.** MicroCT cross section of *Ficus* leaves showing the basic composite structure of cystoliths. (A-D) adaxial cystoliths; (E-H) abaxial cystoliths. (A, E) *F. microcarpa*; (B, F) *F. benjamina*; (C, G) *F lyrata*; (D) *F. benghalensis*; (H) *F. carica*. Scale bars = 60 μm

**Fig. S5.** SEM micrograph of an embedded and polished section through a cystolith of *F. microcarpa*, and table of elemental EDS analyses performed in the designated points: 1) silica stalk; 2, 3) internal ACC phase; 4)bulk external ACC phase

**Fig. S6.** Bar plots of maximum fluorescence intensity from microfluorimetry curves for the 6 *Ficus* species. The data were obtained illuminating (A) adaxially and (B) abaxially. Dark gray bars are for measurements on cystoliths (C) and light gray bars are for measurements on tissue without cystoliths (T). The standard error for each measurement is indicated by the error bar. Leaf cross sections showing cystolith locations and morphologies are schematically represented on the x axes.

**Fig. S7.** Percentage variation of net CO_2_ assimilation for adaxial versus abaxial irradiance. The values are obtained by subtracting to the value measured for each species the value measured for *F. religiosa* (ΔPn_cyst_ (%)). The bar indicates the standard error. Leaf cross sections showing cystolith locations and morphologies are schematically represented on the x axes. Dark gray diamonds: data obtained at 1000 μmol (ph) m^−2^s^−1^ irradiance. Red squares: data obtained at 500 μmol (ph) m^−2^s^−1^ irradiance.

**Fig. S8.** Percentage difference of net CO_2_ assimilation for adaxial versus abaxial irradiance (ΔPn_cyst_ %). Data obtained illuminating at 1000 μmol (ph) m^−2^s^−1^ leaves of greenhouse *Ficus* (dark gray diamonds) and leaves of *Ficus* grown outdoors (red squares). The bar indicates the standard error. For all values above the black line adaxial irradiance is more favorable than in a leaf without adaxial cystoliths, for values below the black line abaxial irradiance is more favorable than in a leaf without abaxial cystoliths. Leaf cross sections showing cystolith locations and morphologies are schematically represented on the x axes. The data show the same trend for outside and greenhouse plants.

**Fig. S9**. Comparison between adaxial-abaxial percent variation in stomatal conductance (blue) and in net carbon assimilation, Pn_ab/ad_% (orange). (A) *F. microcarpa*, (B) *F. benjamina* and (C) *F. lyrata.* Positive values of stomata conductance and Pn show that both values are higher for adaxial irradiance, and negative values show that they are higher for abaxial irradiance.

