## Supplementary material for "Cystoliths in *Ficus* leaves: increasing carbon fixation in saturating light by light scattering off a mineral substrate": Figures S1-S9, Tables S1-S2

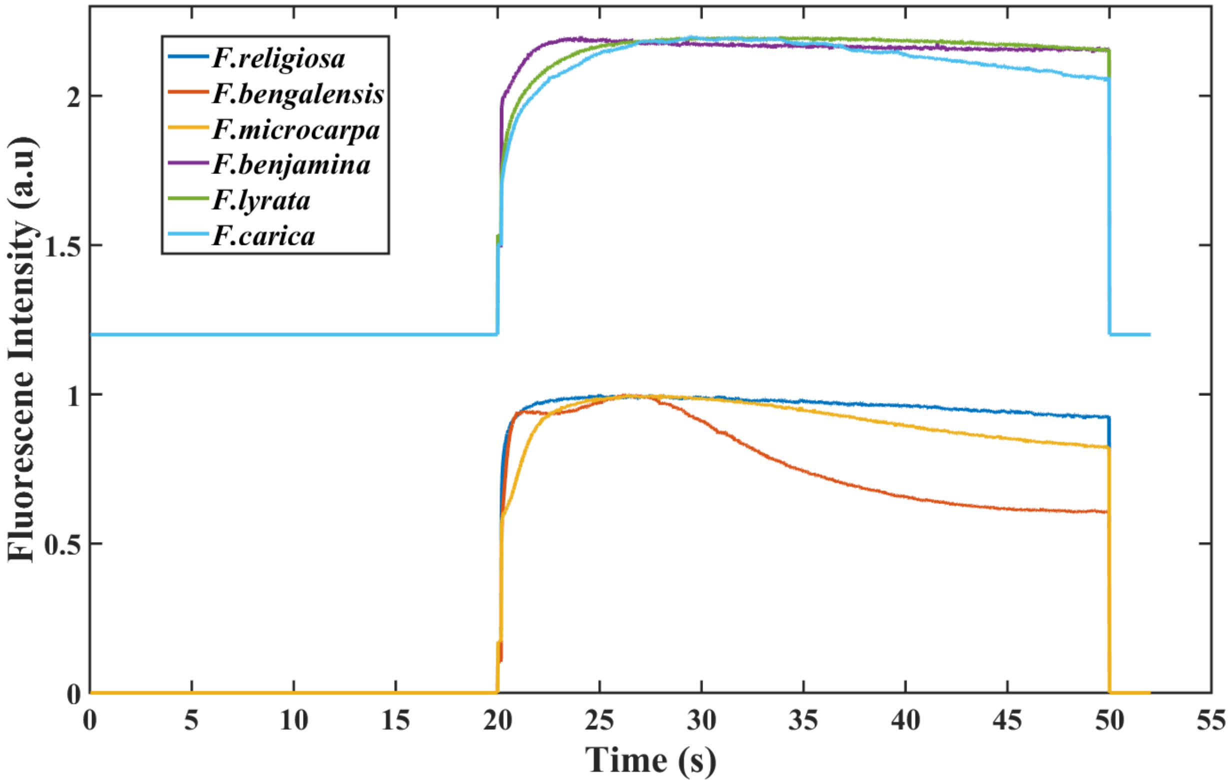


**Figure S1**. Fluorescence intensity evolution over time. The graph shows the dynamics of the measurement for the 6 *Ficus* species on the adaxial side, off crystals. During a 30 s measurement the fluorescence intensity does not varies significantly. The decrease in fluorescence signal is about 10% for all leaves accept *F.bengalensis* with a 40% decrease. The same trend is shown considering adaxial measurements on crystals. From the curves it is also possible to appreciate that the 900 μmol (photon) $m^{-2}s^{-1}$ illumination is saturating all leaves.

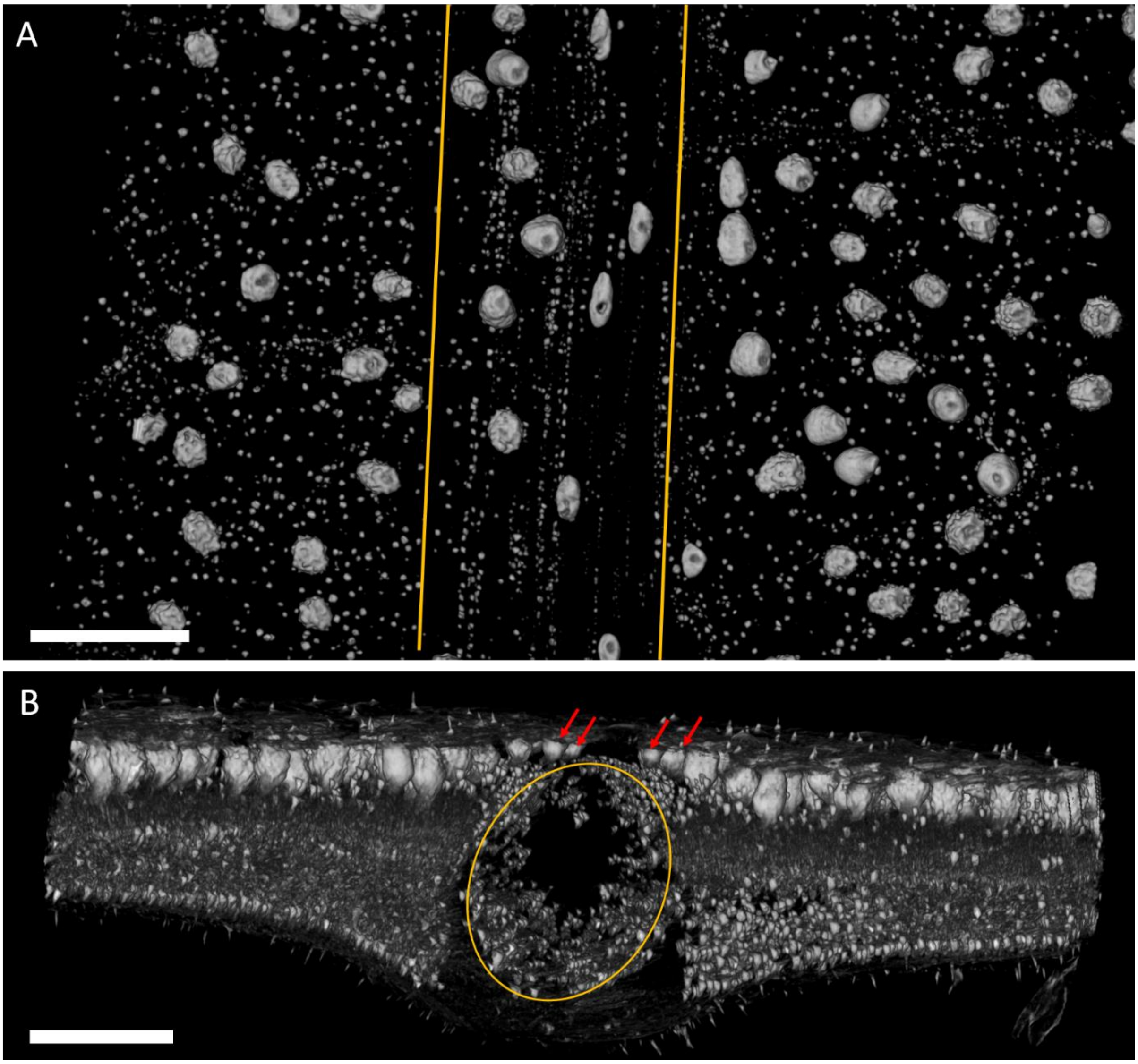


**Figure S2.** MicroCT volumes of *Ficus bengalensis* leaves. A)  top view showing that some of the cystoliths located above the main vein (delimited by yellow lines) are hollow in the center and they are elongated perpendicular to direction of the palisade layer. B) Cross-sectional perspective view showing that some of the cystoliths (red arrows) are deposited also above the veins (the main vein diameter is shown by a yellow circle).

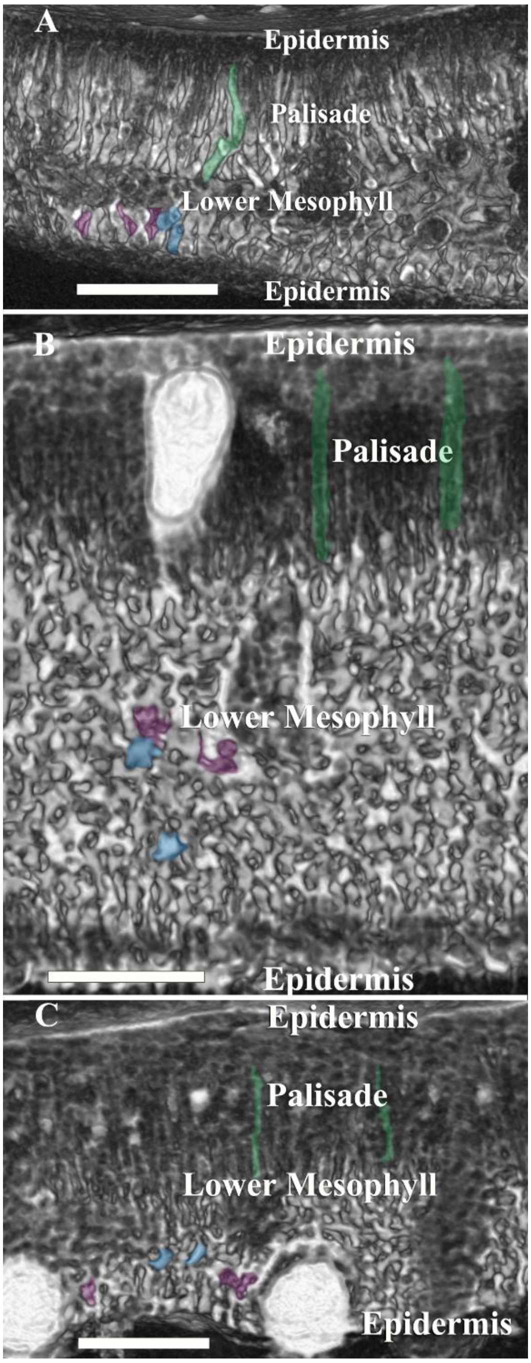


**Figure S3.** MicroCT cross section of *Ficus* leaves showing tissue anatomy. In the palisade and the lower mesophyll single cells are highlighted respectively in green and blue. The air spaces between lower mesophyll cells are purple. (A) *F. religiosa*, (B) *F. lyrata*, (C) *F. carica* . Scale bar 100 µm.

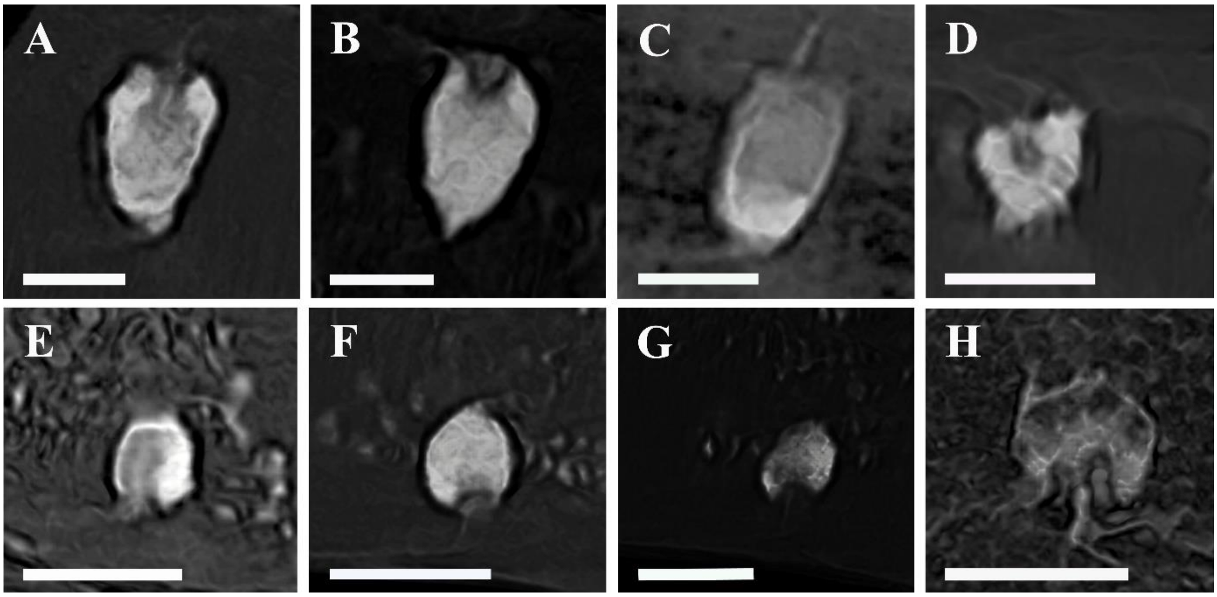


**Figure S4.** MicroCT cross section of *Ficus* leaves showing the basic composite structure of cystoliths. (A-D) adaxial cystoliths; (E-H) abaxial cystoliths. (A, E) *F. microcarpa*; (B, F) *F. benjamina*; (C, G) *F lyrata*; (D) *F. benghalensis*; (H) *F. carica*. Scale bars = 60 µm

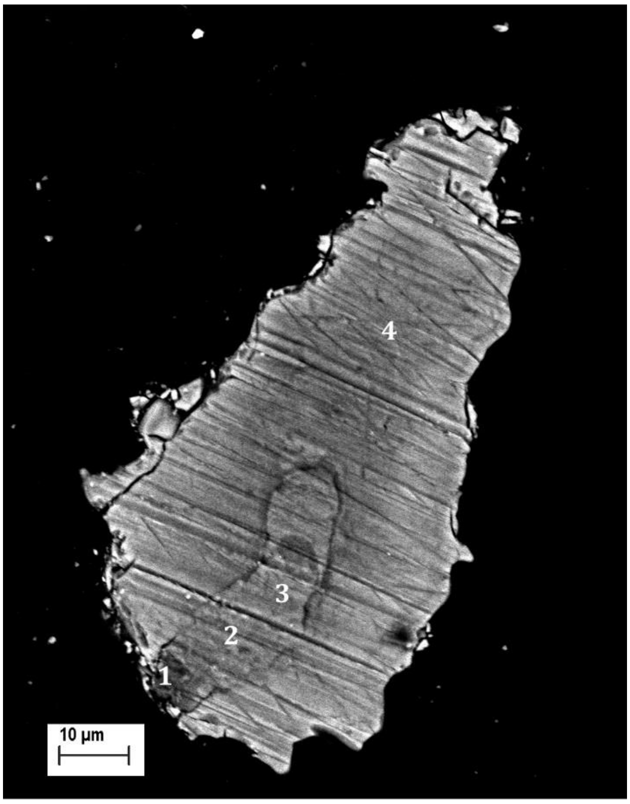


| # | *Si [wt%]* | *Ca [wt%]* | *Mg [wt%]* | *? [wt%]* |
|---|---|---|---|---|
| ***1*** | 48.89 | 22.77 | 19.06 | 9.28 |
| ***2*** | 3.9 | 91.62 | 4.48 | |
| ***3*** | 0.62 | 96.61 | 2.77 | |
| ***4*** | 0.15 | 93.36 | 6.49 | |

**Figure S5.** SEM micrograph of an embedded and polished section through a cystolith of *F. microcarpa*, and table of elemental EDS analyses performed in the designated points: 1) silica stalk; 2, 3) internal ACC phase; 4)bulk external ACC phase

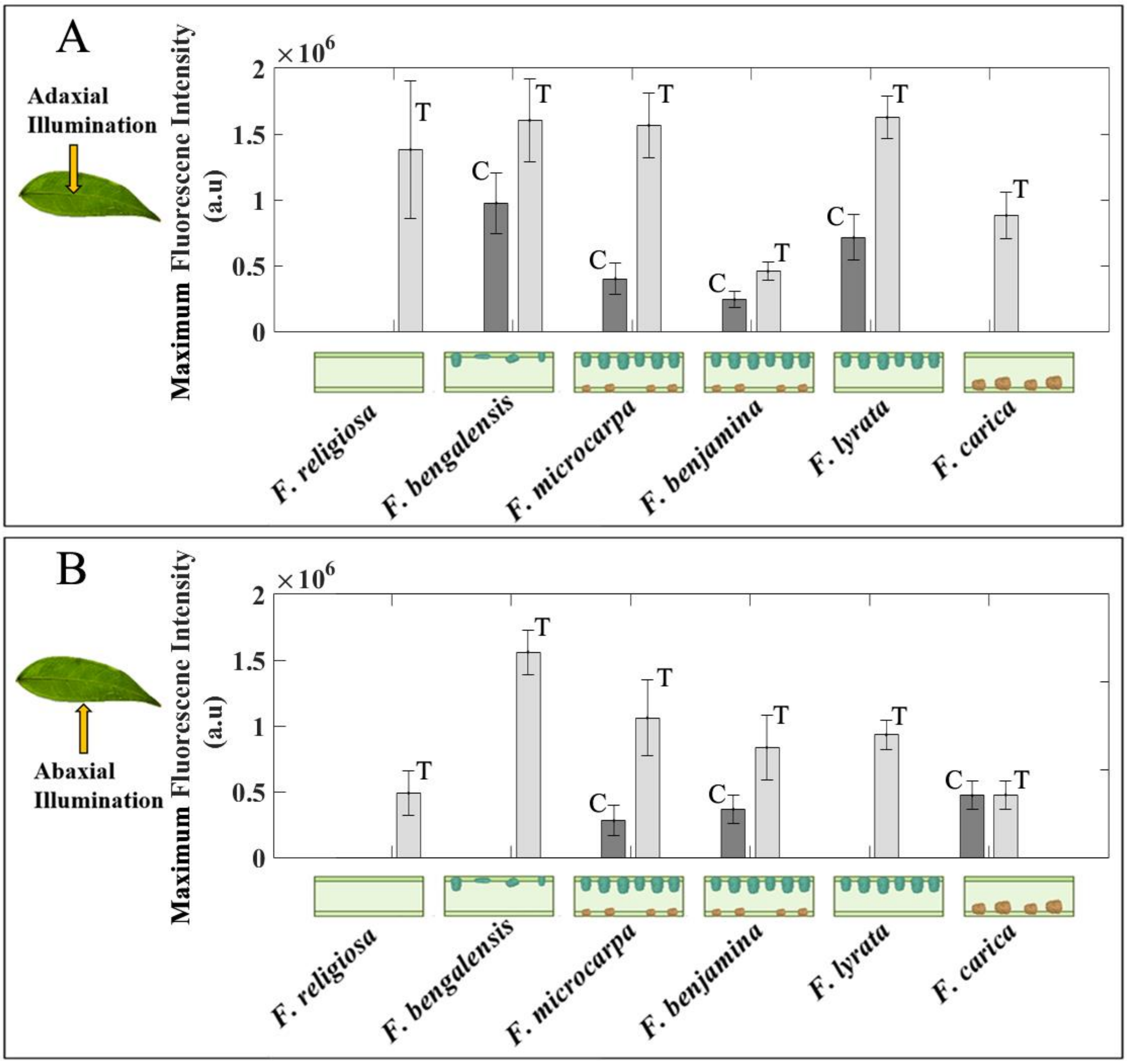


**Figure S6** Bar plots of maximum fluorescence intensity from microfluorimetry curves for the 6 *Ficus* species. The data were obtained illuminating (A) adaxially and (B) abaxially. Dark gray bars are for measurements on cystoliths (C) and light gray bars are for measurements on tissue without cystoliths (T). The standard error for each measurement is indicated by the error bar. Leaf cross sections showing cystolith locations and morphologies are schematically represented on the x axes.

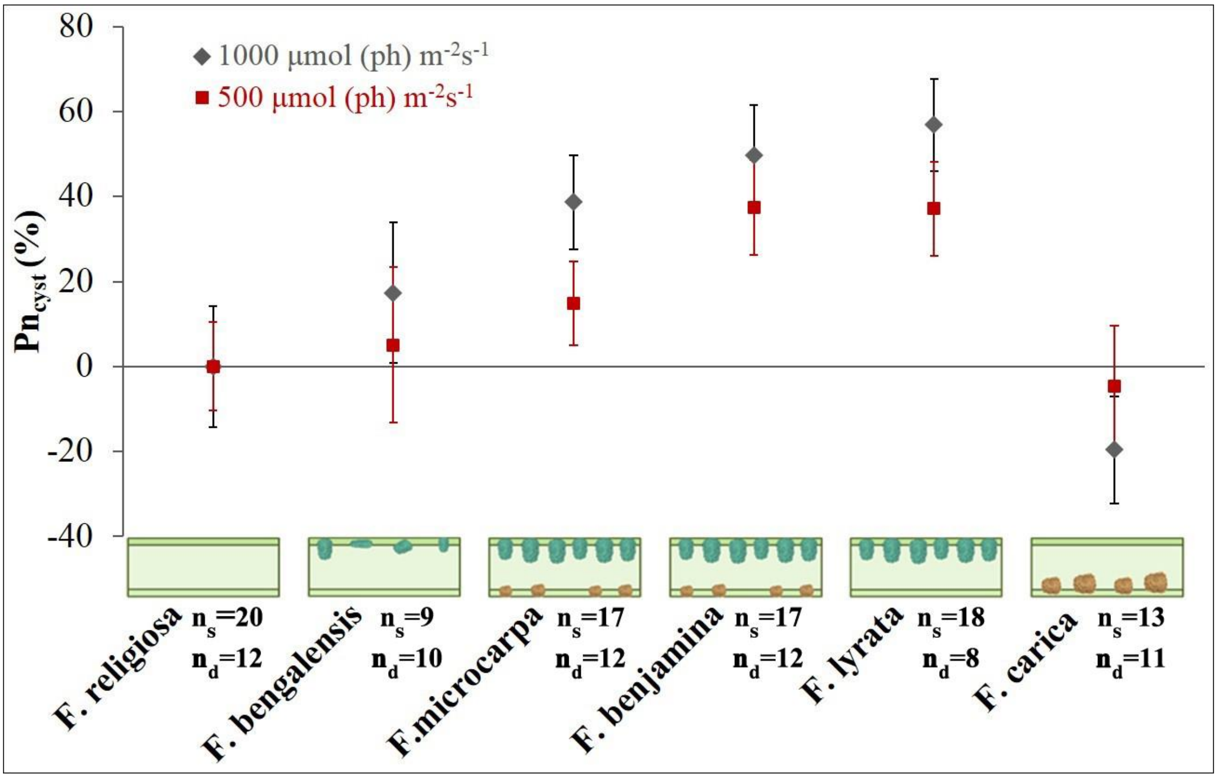


**Figure S7.** Percentage variation of net $CO_2$ assimilation for adaxial versus abaxial illumination. The values are obtained by subtracting to the value measured for each species the value measured for *F. religiosa* (ΔPncyst (%)). The bar indicates the standard error. Leaf cross sections showing cystolith locations and morphologies are schematically represented on the x axes. Dark gray diamonds: data obtained at 1000 µmol (ph) m-2s-1 irradiance ($n_s$ is the number of acquired measurements). Red squares: data obtained at 500 µmol (ph) m-2s-1 irradiance ($n_d$ is the number of acquired measurements).

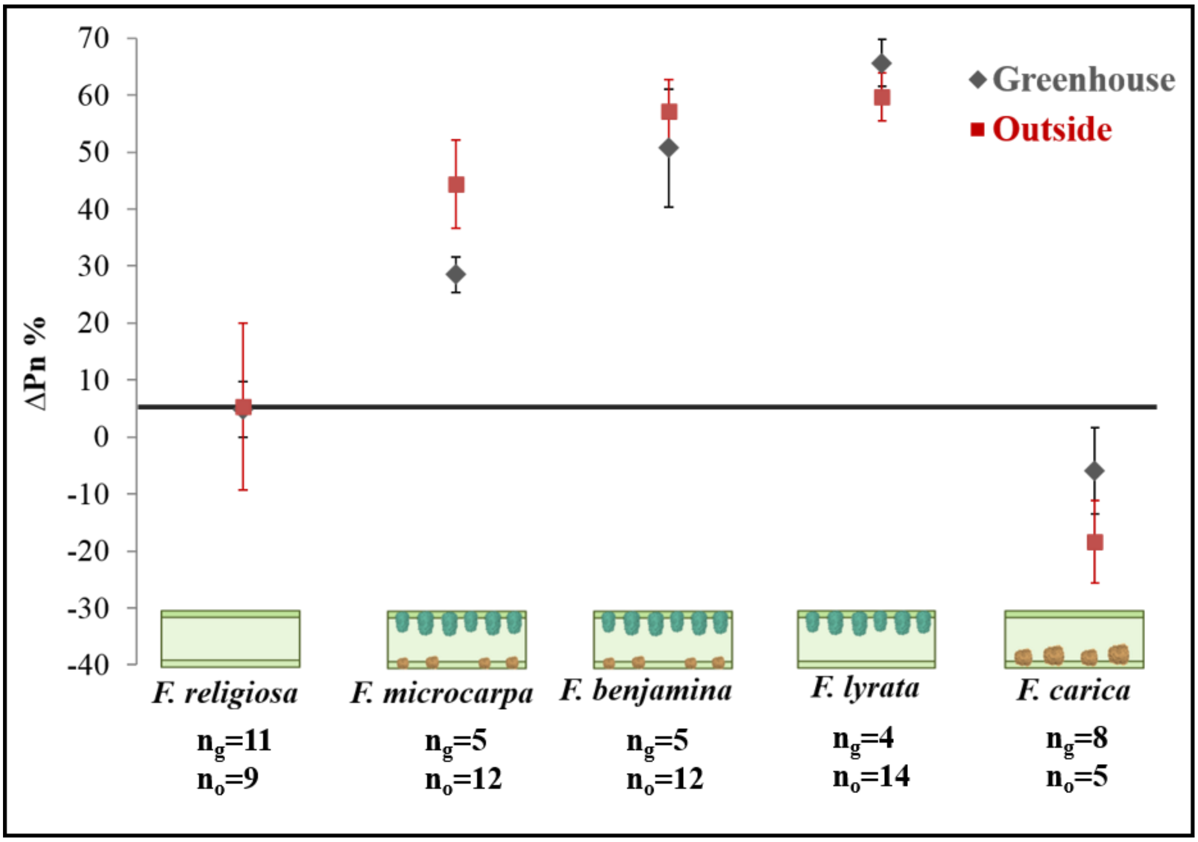


**Figure S8.** Percentage difference of net $CO_2$ assimilation for adaxial versus abaxial illumination (ΔPn%). Data obtained illuminating at 1000 μmol (ph) $m^{-2}s^{-1}$ leaves of greenhouse *Ficus* (dark gray diamonds) and leaves of *Ficus* grown outdoors (red squares). The bar indicates the standard error, $n_g$ is the number of measurements acquired in the greenhouse, $n_o$ is the number of measurements acquired outdoors. For all values above the black line adaxial illumination is more favorable than in a leaf without adaxial cystoliths, for values below the black line abaxial illumination is more favorable than in a leaf without abaxial cystoliths. Leaf cross sections showing cystolith locations and morphologies are schematically represented on the x axes. The data show the same trend for outside and greenhouse plants.

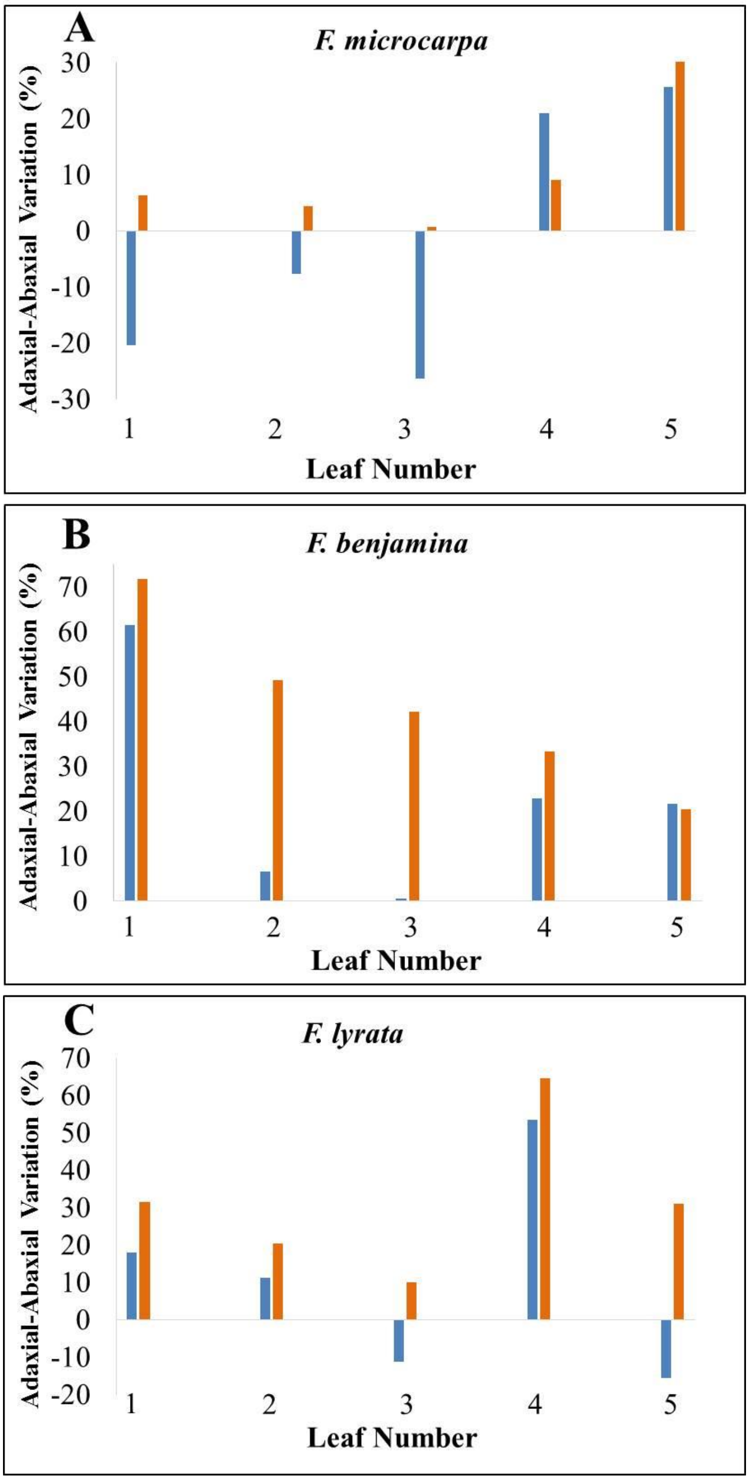


**Figure S9**. Comparison between adaxial-abaxial percent variation in stomatal conductance (blue) and in net carbon assimilation, $Pn_{ab/ad}$% (orange). (A) *F. microcarpa*, (B) *F. benjamina* and (C) *F. lyrata.* Positive values of stomata conductance and Pn show that both values are higher for adaxial illumination, and negative values show that they are higher for abaxial illumination.

**Table S1.** $CO_2$ assimilation measurements were performed on the six Ficus species. For each leaf net $CO_2$ assimilation (Pn) was measured for adaxial illumination (Pn adaxial) and for abaxial illumination (Pn abaxial). The perceptual variation between Pn adaxial and Pn abaxial was calculated (Pn ab-ad% = 100-100*$Pn_{abaxial}$/$Pn_{adaxial}$). P-values from a paired t-test was obtained by testing the difference between $Pn_{adaxial}$ and Pn $_{abaxial}$ The values are indicated in the first column in between brackets. Significant difference at α = 0.05.

| ***F. religiosa*** | **Pn adaxial μmol m$^{-2}$ s$^{-1}$** | **Pn abaxial μmol m$^{-2}$ s$^{-1}$** | **ΔPn ad-ab %** |
|---|---|---|---|
| **Outside Illumination 1000 μmol m$^{-2}$ s$^{-1}$** (P= 0.524) | 0.48 | 0.47 | 1.64 |
| | 2.34 | 2.27 | 2.61 |
| | 2.57 | 2.47 | 4.04 |
| | 0.40 | 0.69 | -41.07 |
| | 0.86 | 1.83 | -52.68 |
| | 4.42 | 1.12 | 74.70 |
| | 2.55 | 3.55 | -28.12 |
| | 1.62 | 0.55 | 65.87 |
| | 1.42 | 1.12 | 21.31 |
| **Greenhouse Illumination 1000 μmol m$^{-2}$ s$^{-1}$** (P= 0.551) | 5.77 | 5.06 | 12.46 |
| | 4.82 | 4.11 | 14.79 |
| | 4.25 | 8.98 | -52.61 |
| | 7.49 | 7.17 | 4.25 |
| | 7.09 | 6.68 | 5.76 |
| | 5.05 | 4.85 | 3.96 |
| | 2.20 | 1.78 | 19.20 |
| | 2.58 | 2.08 | 19.50 |
| | 1.23 | 3.61 | -34.12 |
| | 1.07 | 0.99 | 6.96 |
| | 3.54 | 3.25 | 8.21 |
| **Greenhouse Illumination 500 μmol m$^{-2}$ s$^{-1}$** (P= 0.002) | 5.10 | 4.54 | 11.00 |
| | 5.24 | 5.19 | 0.97 |
| | 6.88 | 6.69 | 2.83 |
| | 6.36 | 5.19 | 18.40 |
| | 5.61 | 5.48 | 2.32 |
| | 6.02 | 5.99 | 0.36 |
| | 1.77 | 1.49 | 15.87 |
| | 3.55 | 1.70 | 52.23 |
| | 3.24 | 2.22 | 31.49 |
| | 3.28 | 2.39 | 27.29 |
| | 3.09 | 1.79 | 42.18 |
| | 3.71 | 2.22 | 40.25 |

| ***F. bengalensis*** | **Pn adaxial μmol m$^{-2}$ s$^{-1}$** | **Pn abaxial μmol m$^{-2}$ s$^{-1}$** | **ΔPn ad-ab %** |
|---|---|---|---|
| **Outside Illumination 1000 μmol m$^{-2}$ s$^{-1}$ (P= 0.039)** | 3.24 | 2.36 | 27.14 |
| | 2.27 | 1.41 | 37.85 |
| | 3.00 | 2.72 | 9.05 |
| | 3.89 | 3.26 | 16.19 |
| | 1.07 | 0.48 | 55.14 |
| | 1.32 | 0.88 | 32.96 |

| | | | |
|---|---|---|---|
| | 3.39 | 0.06 | 18.51 |
| | 2.70 | 2.63 | 2.79 |
| | 2.49 | 2.28 | 8.44 |
| **Outside Illumination 500 µmol $m^{-2}$ $s^{-1}$ (P= 0.068)** | 3.15 | 1.63 | 48.25 |
| | 2.72 | 1.12 | 58.92 |
| | 2.09 | 0.94 | 54.82 |
| | 0.86 | 0.45 | 47.28 |
| | 5.17 | 1.41 | 72.65 |
| | 6.75 | 6.13 | 9.20 |
| | 3.35 | 3.70 | -10.29 |
| | 2.77 | 3.76 | -35.95 |
| | 3.20 | 3.68 | -15.09 |

| ***F. microcarpa*** | **Pn adaxial µmol $m^{-2}$ $s^{-1}$** | **Pn abaxial µmol $m^{-2}$ $s^{-1}$** | **∆Pn ad-ab %** |
|---|---|---|---|
| ***Outside Illumination 1000 µmol $m^{-2}$ $s^{-1}$ (P= 0.0002)*** | *4.01* | *3.39* | *15.42* |
| | *4.67* | *0.99* | *78.90* |
| | *4.60* | *3.18* | *30.73* |
| | *3.66* | *2.05* | *44.14* |
| | *4.19* | *0.96* | *77.15* |
| | *3.74* | *1.30* | *65.24* |
| | *1.94* | *0.88* | *54.52* |
| | *1.62* | *0.47* | *70.78* |
| | *2.39* | *1.39* | *41.81* |
| | *1.29* | *0.52* | *59.48* |
| | *1.47* | *0.80* | *45.31* |
| | *2.66* | *0.98* | *63.20* |
| ***Greenhouse Illumination 1000 µmol $m^{-2}s^{-1}$ (P= 0.002)*** | *7.65* | *5.36* | *29.99* |
| | *7.80* | *4.70* | *39.79* |
| | *6.09* | *4.51* | *25.84* |
| | *8.17* | *6.07* | *25.65* |
| | *6.52* | *5.12* | *21.50* |
| ***Outside Illumination 500 µmol $m^{-2}$ $s^{-1}$ (P= 0.002)*** | *2.61* | *1.00* | *61.92* |
| | *0.34* | *0.19* | *43.01* |
| | *1.26* | *1.02* | *18.76* |
| | *0.82* | *0.78* | *4.31* |
| | *0.98* | *0.57* | *42.38* |
| | *1.28* | *0.77* | *40.27* |
| ***Greenhouse Illumination 500 µmol $m^{-2}$ $s^{-1}$ (P= 0.002)*** | *5.92* | *4.61* | *22.14* |
| | *6.16* | *3.09* | *49.79* |
| | *6.97* | *3.72* | *46.69* |
| | *3.76* | *2.85* | *24.01* |
| | *0.98* | *0.67* | *31.02* |
| | *2.61* | *1.43* | *44.98* |

| *F. benjamina* | Pn adaxial µmol $m^{-2}$ $s^{-1}$ | Pn abaxial µmol $m^{-2}$ $s^{-1}$ | ∆Pn ad-ab % |
|---|---|---|---|
| ***Outside Illumination 1000 µmol*** $m^{-2}$ $s^{-1}$ *(P= 0.001)* | *0.39* | *0.23* | *40.72* |
| | *0.84* | *0.48* | *43.31* |
| | *1.85* | *0.79* | *57.24* |
| | *1.47* | *0.49* | *66.46* |
| | *2.80* | *0.40* | *85.89* |
| | *1.98* | *0.59* | *69.89* |
| | *1.42* | *0.79* | *44.35* |
| | *3.18* | *1.42* | *55.35* |
| | *0.48* | *0.17* | *64.44* |
| | *0.92* | *0.21* | *76.67* |
| | *0.91* | *0.29* | *68.04* |
| | *0.59* | *0.51* | *14.55* |
| ***Greenhouse Illumination 1000 µmol*** $m^{-2}$ $s^{-1}$ *(P= 0.029)* | *0.25* | *0.13* | *45.36* |
| | *0.21* | *0.19* | *12.97* |
| | *1.19* | *0.36* | *69.97* |
| | *3.16* | *1.28* | *59.33* |
| | *3.12* | *1.06* | *66.00* |
| ***Outside Illumination 500 µmol*** $m^{-2}$ $s^{-1}$ *(P= 0.083)* | *0.25* | *0.13* | *45.36* |
| | *0.21* | *0.19* | *12.97* |
| | *1.19* | *0.36* | *69.97* |
| | *3.16* | *1.28* | *59.33* |
| | *3.12* | *1.06* | *66.00* |
| ***Greenhouse Illumination 500 µmol m-2 s-1*** *(P= 0.027)* | *1.37* | *0.17* | *87.79* |
| | *0.47* | *0.13* | *71.76* |
| | *5.37* | *2.69* | *49.80* |
| | *3.86* | *1.00* | *74.08* |
| | *2.43* | *1.32* | *45.95* |
| | *9.66* | *4.52* | *53.19* |
| | *0.39* | *0.23* | *40.72* |

| *F. lyrata* | Pn adaxial µmol $m^{-2}$ $s^{-1}$ | Pn abaxial µmol $m^{-2}$ $s^{-1}$ | ∆Pn ad-ab% |
|---|---|---|---|
| ***Outside Illumination 1000 µmol*** $m^{-2}s^{-1}$ ***(P<0.001)*** | *1.53* | *0.32* | *78.97* |
| | *0.98* | *0.27* | *72.21* |
| | *0.53* | *0.30* | *44.72* |
| | *0.56* | *0.10* | *81.40* |
| | *1.17* | *0.32* | *72.85* |
| | *2.88* | *1.39* | *51.87* |
| | *1.72* | *0.40* | *86.65* |
| | *2.98* | *0.79* | *73.42* |
| | *1.46* | *0.62* | *57.31* |

| | | | |
|---|---|---|---|
| | *2.57* | *0.87* | *66.04* |
| | *1.82* | *0.91* | *49.93* |
| | *1.19* | *0.32* | *73.32* |
| | *0.93* | *0.60* | *35.02* |
| | *1.96* | *0.48* | *75.38* |
| ***Greenhouse Illumination 1000 µmol $m^{-2}s^{-1}$ (P= 0.002)*** | *6.72* | *3.31* | *50.69* |
| | *6.84* | *2.66* | *61.04* |
| | *6.39* | *3.54* | *44.51* |
| | *6.90* | *2.62* | *62.10* |
| ***Greenhouse Illumination 500 µmol $m^{-2}$ $s^{-1}$ (P= 0.001)*** | *3.47* | *2.14* | *38.15* |
| | *2.20* | *1.51* | *31.50* |
| | *4.27* | *1.50* | *64.76* |
| | *2.99* | *2.06* | *31.13* |
| | *5.02* | *3.42* | *31.85* |
| | *1.49* | *0.34* | *77.25* |
| | *2.08* | *0.94* | *54.66* |
| | *1.03* | *0.19* | *81.96* |

| ***F. carica*** | ***Pn adaxial µmol $m^{-2}$ $s^{-1}$*** | ***Pn abaxial µmol $m^{-2}$ $s^{-1}$*** | ***∆Pn ad-ab %*** |
|---|---|---|---|
| *Outside Illumination 1000µmol $m^{-2}$ $s^{-1}$* ***(P= 0.119)*** | *0.80* | *0.88* | *-9.27* |
| | *1.36* | *1.14* | *16.12* |
| | *2.86* | *4.01* | *-40.03* |
| | *4.72* | *6.27* | *-32.91* |
| | *2.93* | *3.19* | *-8.06* |
| | *2.08* | *2.59* | *-24.74* |
| | *1.96* | *2.54* | *-29.67* |
| | *3.48* | *2.98* | *14.37* |
| *Greenhouse Illumination 1000 µmol $m^{-2}s^{-1}$* ***(P=0.889)*** | *3.39* | *4.14* | *-22.24* |
| | *3.66* | *2.36* | *35.46* |
| | *6.60* | *6.25* | *5.33* |
| | *6.57* | *6.02* | *8.35* |
| | *7.41* | *8.53* | *-15.18* |
| *Greenhouse Illumination 500 µmol $m^{-2}$ $s^{-1}$* ***(P= 0.119)*** | *6.58* | *7.21* | *-9.61* |
| | *8.72* | *7.21* | *17.33* |
| | *7.21* | *6.00* | *16.81* |
| | *5.75* | *2.27* | *60.43* |
| | *5.90* | *3.31* | *43.90* |
| | *4.47* | *2.92* | *34.59* |
| | *10.35* | *15.45* | *-49.25* |
| | *10.42* | *7.56* | *27.40* |
| | *9.92* | *7.11* | *28.36* |
| | *3.48* | *3.31* | *4.81* |
| | *4.78* | *4.79* | *-0.08* |

**Table S2.** Average percentage variation between Pn for adaxial illumination and Pn for abaxial illumination ($Pn_{ab/ad}$ (%)) obtained using 1000 µmol $m^{-2}$ $s^{-1}$ and 500 µmol $m^{-2}$ $s^{-1}$ illumination values.

| ***Ficus species*** | **Illumination 1000 µmol $m^{-2}$ $s^{-1}$** | **Illumination 500 µmol $m^{-2}$ $s^{-1}$** |
|---|---|---|
| | **$Pn_{ab/ad}$ (%)** | **$Pn_{ab/ad}$ (%)** |
| ***F. religiosa*** | 6 ± 7 | 20 ± 5 |
| ***F. bengalensis*** | 23 ± 4 | 26 ± 12 |
| ***F.microcarpa*** | 45 ± 4 | 35 ± 5 |
| ***F. benjamina*** | 53 ± 5 | 58 ± 6 |
| ***F. lyrata*** | 63 ± 4 | 58 ± 6 |
| ***F. carica*** | -14 ± 5 | 16 ± 9 |
